## Supplementary Information for "Tracer particles sense local stresses in an evolving multicellular spheroid without affecting the anomalous dynamics of the cancer cells"

(Dated: February 15, 2022)

**Summary:** The Supplementary Information (SI) contains a number of items that are used to support the results in the main text. We first provide the theoretical results based on the numerical solution of the exact coupled stochastic integro-differential (SID) equations, describing the evolution of the inert tracer particles (TP) embedded in a growing multicellular spheroid (MCS). Using the numerical solution for the coupled SID equations and scaling ansatz, we predict the exponents characterizing the dynamics of the TPs in the intermediate ($t \leq \tau$ with $\tau$ being the time in which the CCs divide) and $t \gg \tau$. We then provide additional results from the simulations, which not only validate the theoretical predictions but also elucidate the mechanism that drives the unusual dynamics of TPs, driven by the forces generated by division and apoptosis of the CCs, without affecting the CC dynamics.

**Time-dependent equations for TP and CC densities:** Let us consider tracer particles (TPs) that are embedded in a growing tumor spheroid, as shown Fig. S1a. We model the short-range inter-cell interactions as a sum of two Gaussians that account for repulsion (arising from elastic forces) and attraction (mediated by E-cadherin). The latter accounts for volume excluded from the neighboring TPs and the cancer cells (CCs). In addition, the TPs and CCs are also subject to a random force characterized by a Gaussian white noise spectrum. We assume that the dynamics of the system, consisting of the CCs and TPs (Figure S1 for snapshots generated in simulations) can be described by the over damped Langevin equation,

$$\frac{d\mathbf{r}_i}{dt} = -\sum_{j=1}^{N} \nabla U(|\mathbf{r}_i - \mathbf{r}_j|) + \boldsymbol{\eta}_i(t), \tag{S1}$$

where $\mathbf{r}_i$ is the position of a CC or a TP, and $\boldsymbol{\eta}_i(t)$ is a Gaussian random force with white noise spectrum. The form of $U(|\mathbf{r}_i - \mathbf{r}_j|)$ between a pair of particles (can be either TP-TP, TP-CC or CC-CC) is taken to be ,

$$U(|\mathbf{r}(i) - \mathbf{r}(j)|) = \frac{\nu}{(2\pi\lambda^2)^{3/2}} e^{\frac{-|\mathbf{r}(i)-\mathbf{r}(j)|^2}{2\lambda^2}} - \frac{\kappa}{(2\pi\sigma^2)^{3/2}} e^{\frac{-|\mathbf{r}(i)-\mathbf{r}(j)|^2}{2\sigma^2}}, \tag{S2}$$

where, $\lambda$ and $\sigma$ are the ranges of the repulsive and attractive interactions, and $\nu$ and $\kappa$ are the interaction strengths. Thus, the interactions involving the mixture of CCs and TPs are identical. The potential in Eq.(S2) is one of the model used in the simulations.

When Eq. (S1) is used to describe the TP dynamics, the potential $U_{TP}$ contains both the TP-TP and TP-CC interactions with the corresponding attractive (repulsive) interaction ranges being $\sigma_1(\lambda_1)$ and $\sigma_2(\lambda_2)$, respectively. The potential $U_{CC}$ for the CCs mimics cell-cell adhesion (second term in the above equation), and the excluded volume interactions, and the CC-TP interactions.

A closed form exact equation for the CC density, $\phi(\mathbf{r}, t) = \sum_i \phi_i(\mathbf{r}, t)$ ( $\phi_i(\mathbf{r}, t) = \delta[\mathbf{r} - \mathbf{r_i(t)}]$) can be obtained using the well-known and standard Dean's method [1]. The formally exact time evolution of $\phi(\mathbf{r}, t)$ given by,

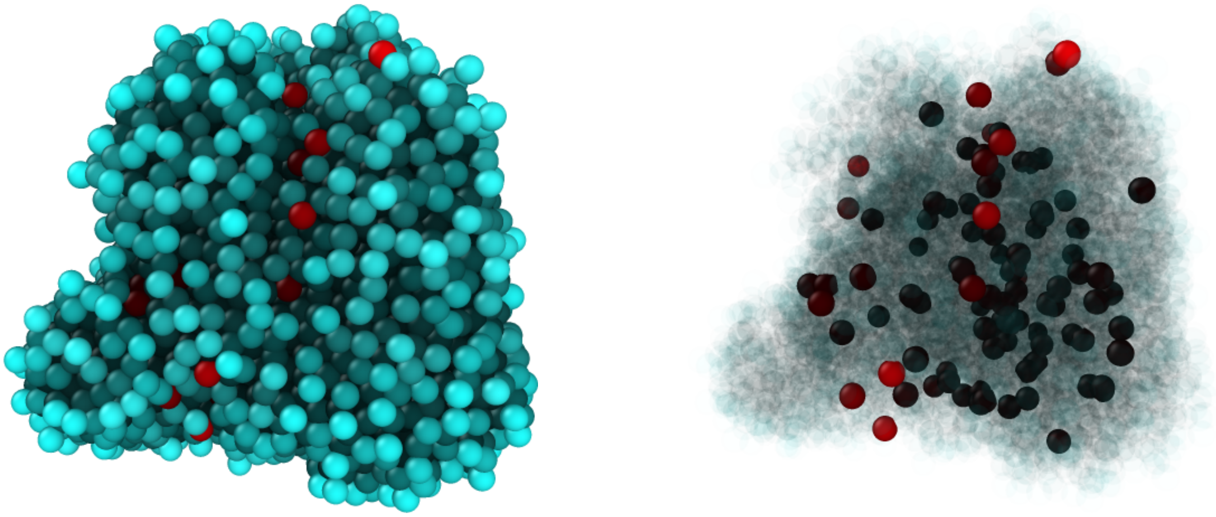

**(a)** **(b)**

**FIG. S1:** Snapshot from tumor simulations with embedded tracers. **(a)** A 3D simulated spheroid consisting of approximately 4800 CCs and 100 TPs. The CCs are in cyan, and the tracers are in red. **(b)** The spheroid was rendered by making the CC cells transparent (light colored cyan) in order to show the interior of the spheroid. The TPs are opaque. Some of the TPs appear black because it is a depiction of a 3D image. The purpose of displaying these snapshots is to visually show that the TPs are randomly distributed within the multicellular spheroid, implying that their migration is largely determined by the forces arising from the CCs.

$$\begin{aligned}\frac{\partial \phi(\mathbf{r},t)}{\partial t} &= D_\phi \nabla^2 \phi(\mathbf{r},t) + \nabla \cdot \Big( \phi(\mathbf{r},t) \int d\mathbf{r}' [\psi(\mathbf{r}',t) \nabla U_{CC-TP}(\mathbf{r}-\mathbf{r}') \\ &\quad + \phi(\mathbf{r}',t) \nabla U_{CC}(\mathbf{r}-\mathbf{r}')] \Big) + \frac{k_a}{2}\phi(\frac{2k_b}{k_a} - \phi) + \nabla \cdot \big( \eta_\phi(\mathbf{r},t) \phi^{1/2}(\mathbf{r},t) \big) + \sqrt{k_b \phi + \frac{k_a}{2}\phi^2} f_\phi \,,\end{aligned} \tag{S3}$$

where $\eta_\phi$ satisfies $< \eta_\phi(\mathbf{r},t)\eta_\phi(\mathbf{r}',t') >= 2D_\phi \delta(\mathbf{r}-\mathbf{r}')\delta(t-t')$ and $f_\phi$ satisfies $< f_\phi(\mathbf{r},t) f_\phi(\mathbf{r}',t') >= \delta(\mathbf{r}-\mathbf{r}')\delta(t-t')$. The term $\propto \phi(\phi_0 - \phi)$ accounts for cell division (rate $= k_b$) and apoptosis (rate $= k_a$), with $\phi_0 = \frac{2k_b}{k_a}$[2, 3]. The coefficient of $f_\phi$, given by $\sqrt{k_b\phi + \frac{k_a}{2}\phi^2}$, is the strength of the noise corresponding to number fluctuations of the CCs, and is a function of the CC density. Similarly, the evolution of the density of TP, $\psi(\mathbf{r},t) = \sum_i \psi_i(\mathbf{r},t) = \sum_i \delta[\mathbf{r} - \mathbf{r_i(t)}]$, may be written as,

$$\begin{aligned}\frac{\partial\psi(\mathbf{r},t)}{\partial t} &= D_\psi\nabla^2\psi(\mathbf{r},t) + \nabla\cdot(\psi(\mathbf{r},t)\int_{\mathbf{r}'}[\psi(\mathbf{r}',t)\nabla U_{TP}(\mathbf{r}-\mathbf{r}') \\ &\quad +\phi(\mathbf{r}',t)\nabla U_{TP-CC}(\mathbf{r}-\mathbf{r}')]) + \nabla\cdot\left(\eta_\psi(\mathbf{r},t)\psi^{1/2}(\mathbf{r},t)\right).\end{aligned} \quad \text{(S4)}$$

where $\eta_\psi$ satisfies $<\eta_\psi(\mathbf{r},t)\eta_\psi(\mathbf{r}',t')>= 2D_\psi\delta(\mathbf{r}-\mathbf{r}')\delta(t-t')$.

The equations for $\phi(\mathbf{r},t)$ and $\psi(\mathbf{r},t)$ constitute coupled non-linear stochastic integro-differential (SID) equations, which are notoriously difficult to solve analytically. Usually approximations, often without establishing their validity, are used to solve the Dean's equation for one component fields. Here, we solve for the first time to our knowledge, by direct numerical integration of the coupled SIDs involving the fields $\phi(\mathbf{r},t)$ and $\psi(\mathbf{r},t)$ (Eqs.(S3) and (S4)). Because the thrust of the theoretical calculations is on the motility of the TPs and how they affect the CC dynamics, we first numerically calculated the correlation functions associated with the TP density field ($\psi(\mathbf{r},t)$). From the decay of the correlation functions and based by the expected scaling behavior, we computed the dynamical exponents for TPs, which is used to characterize the mean-square displacement (MSD $\sim t^{2/z}$), with $z$ being the dynamic exponent.

By numerically integrating Eqs. S3 and S4, we calculated the density-density correlations for the TPs, $C_{\psi\psi}(t) = \int d^3\mathbf{r} C_{\psi\psi}(\mathbf{r},t)$ where $C_{\psi\psi}(\mathbf{r},t) =<\psi(\mathbf{r},t)\psi(\mathbf{r},t)> - <\psi(\mathbf{r},t)><\psi(\mathbf{r},t)>$. Based on the earlier works [4–6] we expect that the decay of TP density correlation function $C_{\psi\psi}(t)$ should decays as $C_{\psi\psi}(t) \sim t^{1-\frac{2}{z}-\frac{d}{z}}$ ($d$ is the spatial dimension). By fitting the expected scaling behavior for $C_{\psi\psi}(t)$ to the numerical solution to the SIDs the value of $z$ can be extracted. The MSD exponents are given by $2/z$.

Using the numerical results we calculated $C_{\psi\psi}(t)$ shown in Fig.(2a) in the main text. We find that both at short ($t < \tau$) and long times ($t > \tau$) $C_{\psi\psi}(t)$ decays as a power law. By fitting the short time decay to $t^{1-2/z-d/z} \approx t^{-3/7}$ (green dashed line in Fig.(2a) in the main text) we extract $z = 7/2$. The corresponding MSD exponent $\beta_{TP} = 2/z = 4/7$, which implies that at $t/\tau$ less than unity the TP motion is sub-diffusive. A similarly, at $t/\tau > 1$, we find that $z = 7/8$, which leads to the MSD exponent $\alpha_{TP} = 16/7$. Thus, the TP dynamics changes sub-diffusive (glass-like) behavior at short times to hyper-diffusive at long times. The crossover occurs when the cell proliferation becomes relevant. A summary of the values of the exponent is given in Table II.

**Control simulation:** As stated in the main text, TPs, regardless of their nature would be useful as local stress sensor only if they do not significantly alter the dynamics or the local stresses of the CCs. To ascertain if this holds we performed a variety of simulations by changing the TP size, the cell division time, and turning off the interaction between the TPs. The results of these simulations are described below.

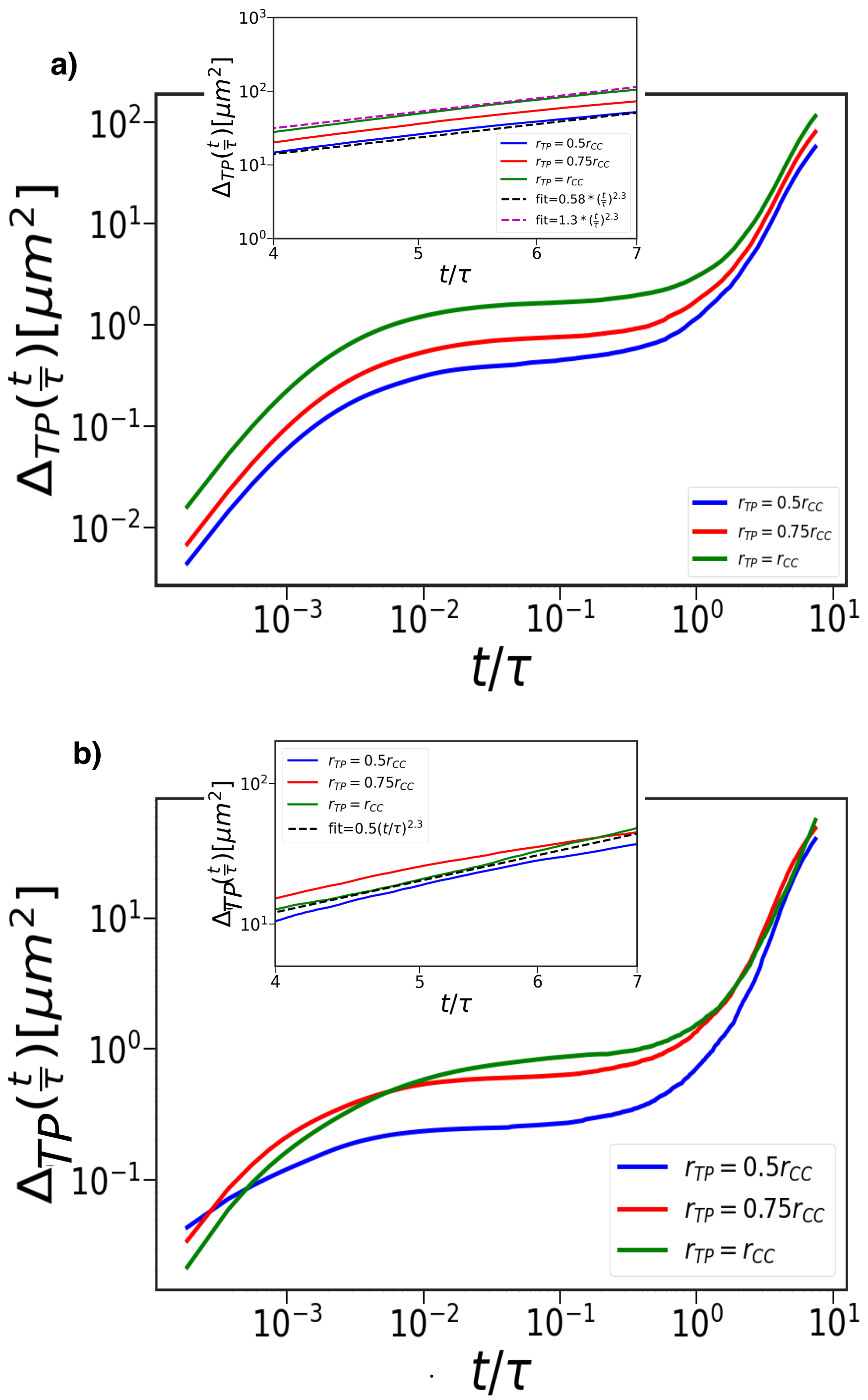


**FIG. S2:** Dependence of the TP size on $\Delta_{TP}$ as a function of $t/\tau$. **(a)** Data are for the Hertz potential (see equations (3)-(5) in the main text). From top to bottom, the curves correspond to decreasing TP radius ($r_{TP} = r_{CC}$ (green), $r_{TP} = 0.75r_{CC}$ (red) and $r_{TP} = 0.5r_{CC}$ (blue), where $r_{CC} = 4.5\mu m$ is average CC radius). TPs with larger radius have larger MSD values in the intermediate time ($\frac{t}{\tau} \leq \mathcal{O}(1)$). In the inset, we focus on the hyper-diffusive regime. The black and magenta dashed line serves as a guide to the eye with $\alpha_{TP} = 2.3$ **(b)** Same as (a) but the results are for the Gaussian interactions.

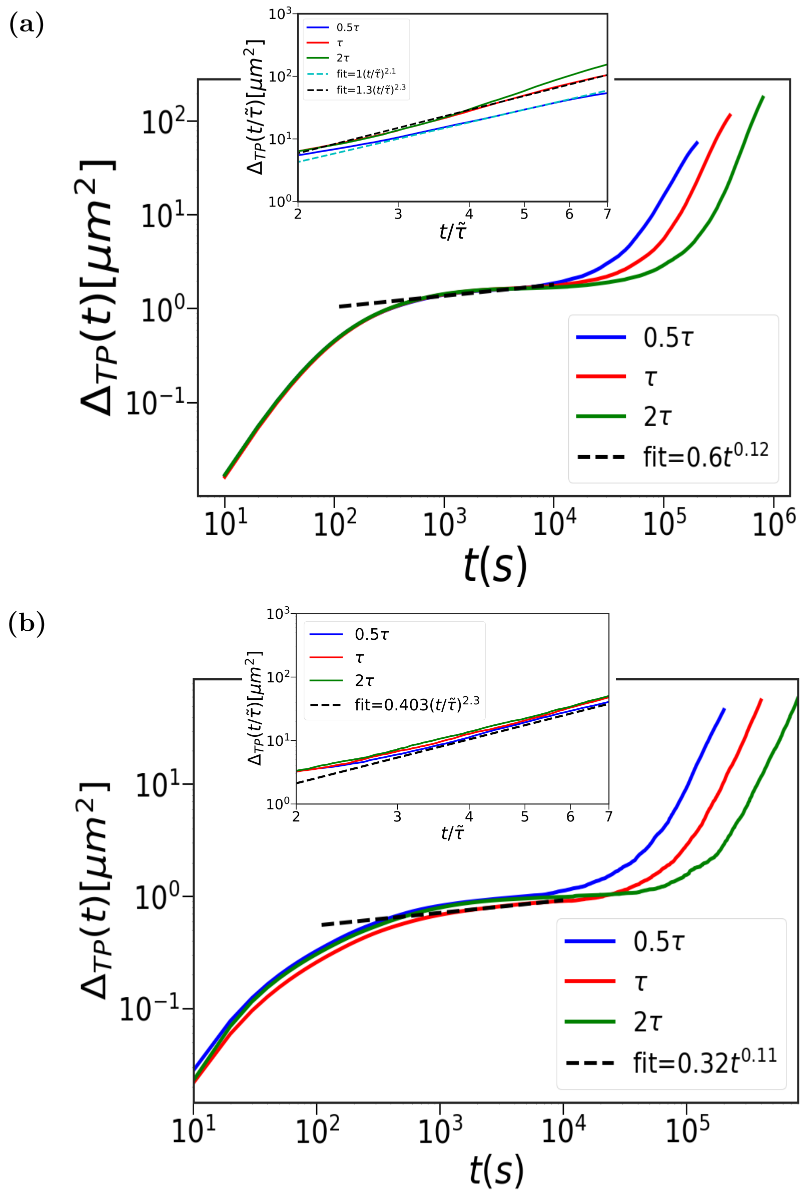


**FIG. S3:** Impact of cell division time $\tau$ on $\Delta_{TP}(t)$ as a function of time $(t)$ for the two types of CC interactions. **(a)** $\Delta_{TP}$, calculated using Eq.(5) in the main text, for the forces between the CCs and TPs. The curves are for 3 cell cycle times (blue – $0.5\tau$, red – $\tau$, and green – $2\tau$). Time taken to reach the super-diffusive limit, which is preceded by a sub-diffusive (glass-like or jammed) regime, increases as $\tau$ increases. In the long-time, $\Delta_{TP}(t)$ exhibits hyper-diffusive motion ($\Delta_{TP} \sim t^{\alpha_{TP}}$ with $\alpha_{TP} > 2$), which is shown in the inset for three $\tau$ values. The $x$-axis in the inset plot is scaled by $\frac{1}{\tilde{\tau}}$. The values of $\tilde{\tau}$ are $0.5\tau$, $\tau$, and $2\tau$. The black (cyan) dashed line shows exponent $\alpha_{TP} = 2.3$ (2.1). The curve with 0.5 $\tau$ is best fit using $\alpha_{TP} = 2.1$. **(b)** Same as (a) except the Gaussian potential (Eq.(6) in the main text) is used in the simulations. Interestingly, $\alpha_{TP}$ does not change appreciably, implying that the long time dynamics is impervious to changes in the short range systematic interactions.

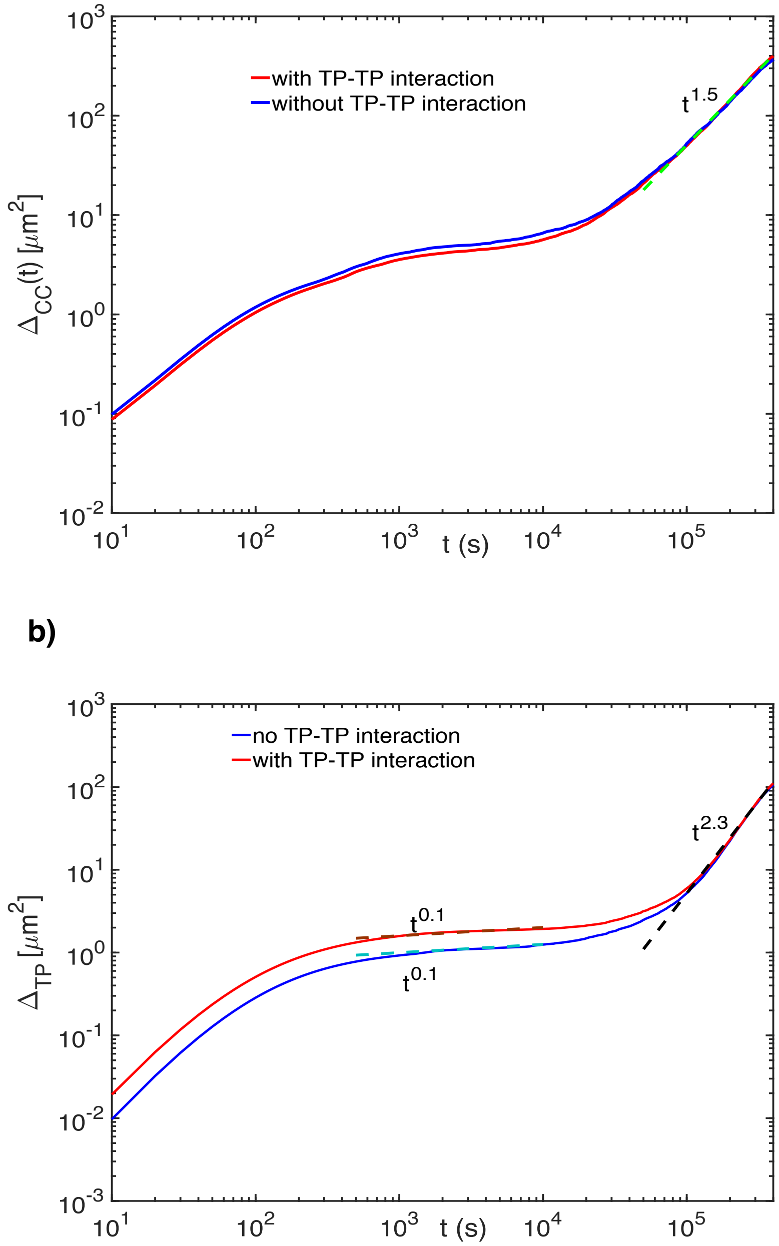


.

**FIG. S4: TP-TP interactions do not affect the long time dynamics of the TPs or CCs.** (a) $\Delta_{CC}$ with (red curve) and without (blue) TP-TP interactions. The TP-TP interactions play no role in the CC dynamics. The cyan dashed line shows $\alpha_{CC} = 1.5$ for both the cases. (b) $\Delta_{TP}$ with (red curve) and without (blue) TP-TP interaction. $\Delta_{TP}$ differs in magnitude at intermediate times. However, TP-TP interactions do not impact the long time hyper-diffusive dynamics of the TPs.

| Definition | Parameters | Value |
|---|---|---|
| Cell diffusion constant | $D_\phi$ | $10^{-8}\mu m^2/s$ |
| Tracer diffusion constant | $D_\psi$ | $10^{-8}\mu m^2/s$ |
| Cell division rate | $k_b$ | $1/\tau,\ \tau = 54,000s$ |
| Cell apoptosis rate | $k_a$ | $0.1k_b$ |
| Box size | $L$ | $500\ \mu m$ |
| Repulsive interaction range | $\lambda$ | $10\ \mu m$ |
| Attractive interaction range | $\sigma$ | $10\ \mu m$ |
| Repulsive interaction strength | $\nu$ | $103\mu N \cdot \mu m^4$ |
| Attractive interaction strength | $\kappa$ | $100\mu N \cdot \mu m^4$ |
| Integration time step | $\delta t$ | $0.01\tau$ |
| Poisson ratio | $\nu_i$ | 0.5 |
| Elastic modulus | $E_i$ | $10^{-3}MPa$ |
| Adhesive coefficient | $f^{ad}$ | $10^{-4}\mu N/\mu m^2$ |

**TABLE I:** Parameters (except last three) used in the numerical integration of Eqs. S3 and S4. The values of the additional parameters (last three) used in the simulations.

$\alpha_{TP}$ **is nearly independent of the TP size:** We varied the radius of the TP ($r_{TP}$) from $0.5r_c$ to $r_c$, where $r_c = 4.5\mu m$ is the average CC radius. Figure S2 shows $\Delta_{TP}(t)$ as function of $t$ for the Hertz potential (Eq.(5) in the main text). Similar behavior is found for the Gaussian potential as well. In the intermediate time regime, larger sized TPs have higher MSD because they experience greater repulsive forces due to bigger excluded volume interactions. In the long time limit, $\Delta_{TP}$ exhibits hyper-diffusion (insets in Figure S2). The CC-TP interaction term, $\nabla \cdot \big(\psi_1(\mathbf{r},t)\int d\mathbf{r}'\phi_1(\mathbf{r}',t)\nabla U(\mathbf{r}-\mathbf{r}')\big)$, in Eq.(S4) shows that the radius merely alters the interaction strength, and does not fundamentally change the scaling behavior. The conclusion that the values of $\alpha_{TP}$ do not change, anticipated on theoretical grounds, is supported by simulations.

**CC cell division does not significantly alter the TP dynamics:** Fig.(S3) shows the influence of cell division on $\Delta_{TP}(t)$ for the Hertz and Gaussian potentials, described in the Methods section in the main text. It is clear that by varying cell division time from $(0.5-2)\tau$ the short time glassy dynamics as well as the hyper-diffusive behavior are maintained. Even the values of the exponents ($\beta_{TP}$ and $\alpha_{TP}$) are not significantly altered.

**Effect of TP-TP interactions:** Typically, in experiments the number of TPs in a given MCS is only about four or five. It is unlikely there is any interaction between the TPs. Even though we use 100 TPs they are sufficiently far away that $U_{TP-TP} \approx 0$. We performed simulations with and without TP-TP interactions. Fig.(S4) clearly shows that

(a)

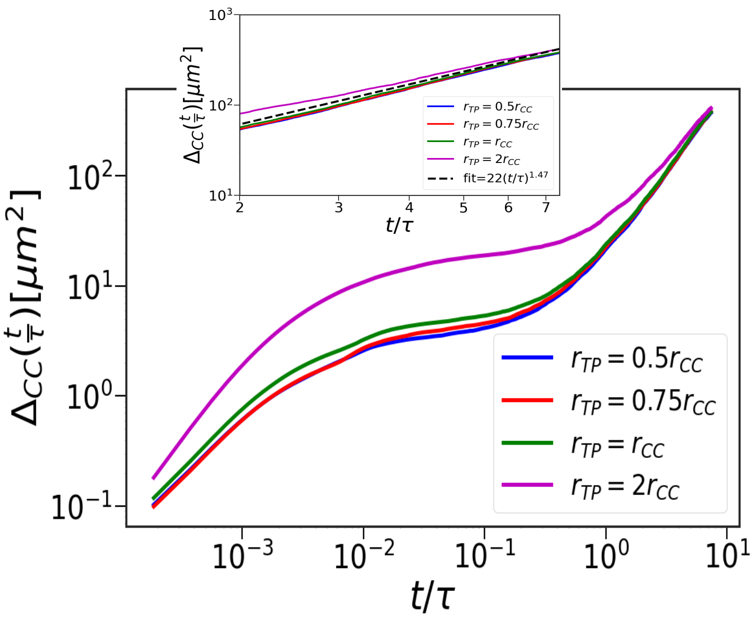


(b)

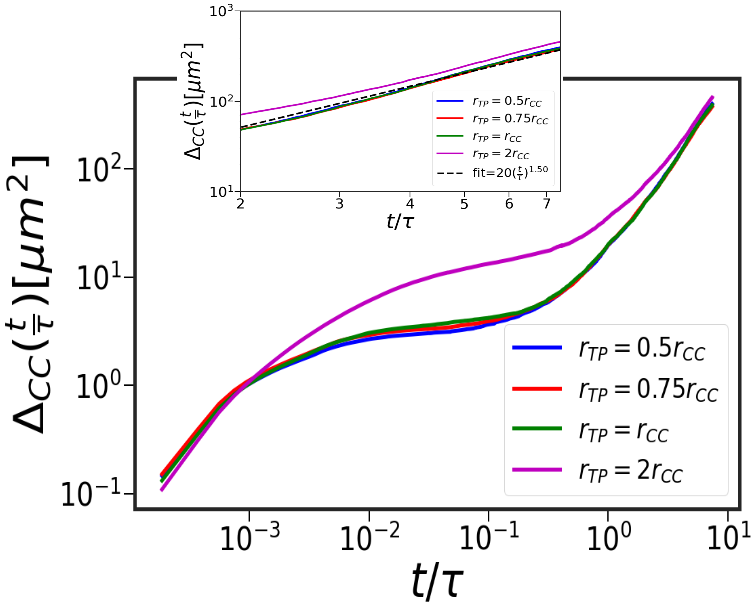


**FIG. S5:** Influence of TPs on CC dynamics for two cell-cell potentials. (a) $\Delta_{CC}$, as function of $t/\tau$, using the Hertz potential (Eq.(5) in the main text). From top to bottom, the curves are for different values of the radius of the TPs (magenta $r_{TP} = 2r_{CC}$, green $r_{TP} = r_{CC}$, red $r_{TP} = 0.75r_{CC}$ and blue $r_{TP} = 0.5r_{CC}$ (appears to be hidden), where $r_{CC} = 4.5\ \mu m$ is the average cell radius). In the intermediate times, $\Delta_{CC}(t)$ is larger for TPs with larger radius. The inset focuses on long times ($\frac{t}{\tau} > 1$). The black line is a guide to show the value of $\alpha_{CC} = 1.47$. (b) Same as (a) but with the Gaussian potential. It is note worthy that the long time MSD exponent for the CCs is unaffected by the inert TPs, even after seven cell divisions.

| | Theory | Hertz | Gaussian |
|---|---|---|---|
| $\beta_{TP}$ | 0.57 | 0.12 | 0.11 |
| $\alpha_{TP}$ | 2.28 | 2.30 | 2.30 |
| $\alpha_{CC}$ | 1.45 | 1.47 | 1.50 |

**TABLE II:** MSD exponents from theory and simulations.

neither $\Delta_{TP}(t)$ nor $\Delta_{CC}(t)$ is affected by TP-TP interactions.

**Influence of the TPs on the CC dynamics:** We calculated $\Delta_{CC}(t)$ as a function of $t$ using the Hertz (Gaussian) potential as a function of the TP radius (Figure S5**a** (S5**b**)). For both the potentials, the values of $\Delta_{CC}(t)$ for $t < \tau$ increases as the TP sizes increase. Before cell division, the number of CCs and TPs are similar, which explains the modest influence of the TPs on the dynamics of the CCs in the intermediate time regime. The larger TPs experience stronger repulsion (the repulsive interaction is proportional to $R^2$) initially, which increases the magnitude of $\Delta_{CC}(t)$. The long-time dynamics is not significantly affected by the CC-TP interactions (see Figure S5). In the absence of the TPs, the CCs exhibit super-diffusion where the MSD scales as $t^{\alpha_{CC}}$ with $\alpha_{CC} = 1.33$[7]. In the presence of the TPs, the CC dynamics remains super-diffusive with $\alpha_{CC} \approx 1.45$, which shows that the TPs do not alter the CC dynamics significantly. The results in figures (S2)-(S5) show that the influence of TPs on the CCs is minimal, which fully justifies their use as pressure sensors. The predictions for the CC dynamics made here remains to be tested.

**Straigtness Index:** The Straightness Index (SI) is, given by, $SI_i = \frac{\mathbf{r}_i(t_f)-\mathbf{r}_i(0)}{\sum \delta\mathbf{r}_i(t)}$, where $\mathbf{r}_i(t_f) - \mathbf{r}_i(0)$ is the net displacement of $i^{th}$ TP or CC. The denominator, $\sum \delta\mathbf{r}_i(t)$, is the total distance traversed. Fig. S6 shows that the TP trajectories are more straight (higher SI values on an average) or persistent over longer duration. During each cell division, the CCs are placed randomly causing the trajectories to be less persistent, thus explaining the decreased persistence in their motion. In contrast, the TPs experience an impulsive kick in the vicinity of the CCs that undergo cell division. The kick results in a force on the TPs, which tends to produce persistent motion, thus giving rise to the higher SI values, as shown in Fig.(S6).

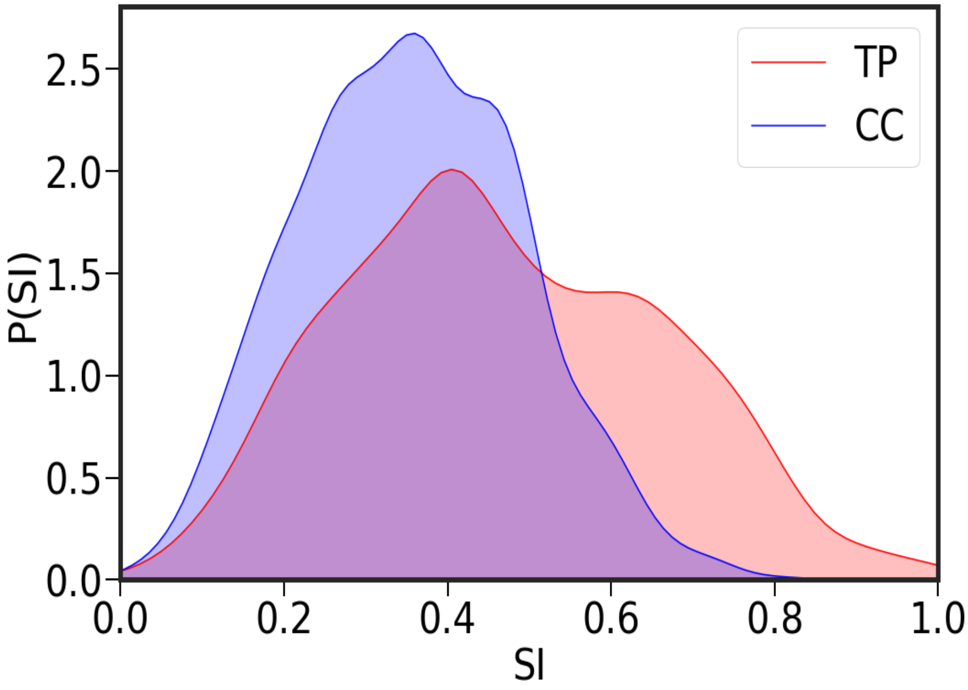


**FIG. S6:** Distribution of the Straightness Index (SI). The red (blue) plot for the TPs (CCs) shows that the TP trajectories are more rectilinear than the CCs.
